# Deciphering cattle temperament measures derived from a four-platform standing scale using genetic factor analytic modeling

**DOI:** 10.1101/2020.01.20.913343

**Authors:** Haipeng Yu, Gota Morota, Elfren F. Celestino, Carl R. Dahlen, Sarah A. Wagner, David G. Riley, Lauren L. Hulsman Hanna

**Author notes:** Corresponding authors: Lauren L. Hulsman Hanna, Department of Animal Sciences, North Dakota State University, DEPT 7630, P.O. Box 6050, Fargo, North Dakota 58108 USA., Gota Morota, Department of Animal and Poultry Sciences, Virginia Polytechnic Institute and State University, 175 West Campus Drive, Blacksburg, Virginia 24061 USA.

## Abstract

The animal’s reaction to human handling (i.e., temperament) is critical for work safety, productivity, and welfare. Subjective phenotyping methods have been traditionally used in beef cattle production. Even so, subjective scales rely on the evaluator’s knowledge and interpretation of temperament, which may require substantial experience. Selection based on such subjective scores may not precisely change temperament preferences in cattle. The objectives of this study were to investigate the underlying genetic interrelationships among temperament measurements using genetic factor analytic modeling and validate a movement-based objective method (four-platform standing scale, FPSS) as a measure of temperament. Relationships among subjective methods of docility score (DS), temperament score (TS), 12 qualitative behavior assessment (QBA) attributes and objective FPSS including the standard deviation of total weight on FPSS over time (SSD) and coefficient of variation of SSD (CVSSD) were investigated using 1,528 calves at weaning age. An exploratory factor analysis (EFA) identified two latent variables account for TS and 12 QBA attributes, termed *difficult* and *easy* from their characteristics. Inclusion of DS in EFA was not a good fit because it was evaluated under restraint and other measures were not. A Bayesian confirmatory factor analysis inferred the *difficult* and *easy* scores discovered in EFA. This was followed by fitting a pedigree-based Bayesian multi-trait model to characterize the genetic interrelationships among *difficult*, *easy*, DS, SSD, and CVSSD. Estimates of heritability ranged from 0.18 to 0.4 with the posterior standard deviation averaging 0.06. The factors of *difficult* and *easy* exhibited a large negative genetic correlation of −0.92. Moderate genetic correlation was found between DS and *difficult* (0.36), *easy* (−0.31), SSD (0.42), and CVSSD (0.34) as well as FPSS with *difficult* (CVSSD: 0.35; SSD: 0.42) and *easy* (CVSSD: −0.35; SSD: −0.4). Correlation coefficients indicate selection could be performed with either and have similar outcomes. We contend that genetic factor analytic modeling provided a new approach to unravel the complexity of animal behaviors and FPSS-like measures could increase the efficiency of genetic selection by providing automatic, objective, and consistent phenotyping measures that could be an alternative of DS, which has been widely used in beef production.

## Introduction

Temperament in cattle traditionally refers to the animal’s behavior in the bail (Tulloh, 1961) or the reaction of animal to human handling (Burrow and Dillon, 1997). Previous studies have revealed cattle temperament has a significant relationship with production, reproduction, immunity, and carcass traits (King et al., 2006; Burdick et al., 2011; Haskell et al., 2014). Additionally, temperament is usually evaluated at an earlier stage of cattle than some production traits. Therefore, temperament could be considered as an indicator of production traits in genetic selection, where selection on temperament can provide an opportunity to improve production and efficiency in the beef industry. Temperament is a complex trait that comprises various behavioral characteristics such as shyness-boldness, exploration avoidance, activity, sociability, and aggressiveness (Réale et al., 2007). Several subjective methods were proposed to score temperament, including temperament scoring of cattle handled in a crush with head bail (Tulloh, 1961), flight distance (Fordyce et al., 1982), docility test (Le Neindre et al., 1995), chute test (Tier et al., 2001), race score (Turner et al., 2011) and qualitative behavior assessment (QBA) (Sant’Anna and da Costa, 2013). These subjective methods are able to integrate various levels of temperament attributes (e.g., calmness, agitation, flightiness, agressiveness) into a single score and create a standardized test by taking advantage of the experience and interpretation of the human evaluator on cattle. This is advantageous for typical production operations due to ease of capturing data compared to objective methods that require specialized equipment (e.g., exit velocity). Even so, closely working with cattle may cause potential danger for evaluators during scoring. Furthermore, there is a concern with evaluation bias in subjective methods, which makes comparison of temperament scoring methods across experiments difficult. Due to this, measurements without human interpretation, such as exit velocity (Burrow et al., 1988), movement-measuring-devices (Stookey et al., 1994; Sebastian et al., 2011), strain gauges (Schwartzkopf-Genswein et al., 1997), and objective chute score (Bruno et al., 2018), have been tested to provide objective and quantifiable temperament measurements. Understanding how these objective measures relate to behavioral attributes is of interest, where most studies have only compared a few common subjective methods with objective methods using a standard multi-trait model (Burrow et al., 1988; Stookey et al., 1994; Sebastian et al., 2011; Bruno et al., 2018). Computational limitations have also hindered further research in understanding relationship between objective and subjective methods of temperament. Therefore, this study introduces a novel, potentially cost effective objective method using a four-platform standing scale (FPSS) and investigates its relationship with subjective methods of docility score (DS), temperament score (TS), and qualitative behavior assessment (QBA) attributes. The objectives of this study were to apply genetic factor analytic modeling to characterize the underlying genetic interrelationships among temperament measures and validate FPSS as an objective measurement of cattle temperament. We employed new statistical approaches, explanatory factor analysis (EFA) and confirmatory factor analysis (CFA), to overcome the computational challenges due to a large number of correlated subjective measurements. To our knowledge, this is the first study to investigate the novel objective measurement FPSS using EFA and CFA for the genetic analysis of cattle temperament.

## Materials and methods

### Animals

From 2014 to 2017, data were collected at weaning time (late September to late October) at North Dakota State University Central Grasslands Research Extension Center (CGREC) near Streeter, North Dakota. Calves (*n* = 1, 528), including 749 heifers and 779 steers, were scored at the weaning age (average age was 161.0 +/− 17.0 d) and included for analysis. Calves were either sired by Angus bulls (all 4 yr) or by Hereford bulls (3 yr) and were from Angus-influenced (all 4 yr) or Hereford by Angus-influenced (2 yr) dams. Calves were assigned to one of two primary breeds (50% or greater) based on known breed percentages, which resulted in 1,340 Angus-based (666 heifers, 674 steers) and 188 Hereford-based (83 heifers, 105 steers) calves. A pedigree including 109,703 animals was formed using the information of dams and records of complete ancestry for registered bulls provided by the American Angus Association and American Hereford Association. All procedures involved in data collection were reviewed and approved by the Institutional Animal Care and Use Committee of North Dakota State University.

### Experiment procedure

The details of temperament evaluation procedure during the first year of data collection was previously described by Hulsman Hanna et al. (2019) and was repeated the remaining 3 years of the study except for collection of the blood draw effect. Briefly, calves were moved through the working pens to the evaluation areas and then sorted to different holding pens for management. In a given year, four evaluators were randomly assigned two of three subjective scoring methods (DS, TS, and QBA) prior to evaluation. A total of 11 evaluators were presented over the 4 yr period as some evaluators were not able to return for all years of the study. Averages per animal for each subjective method that had at least three evaluator scores were used in this study since the focus of this study was not evaluator variation. Basic means and standard deviations are presented in Supplementary Table S1. The first method scored was DS, which is a six point scale where one and six refer to calm and aggressive, respectively (Beef Improvement Federation, 2018). The evaluation of DS was done at the silencer chute with the head of the calf caught and each calf was evaluated less than one minute. Following the evaluation of DS, weaning weight of the calf was recorded when its body was squeezed. Upon released from silencer chute, the calf then entered the FPSS (Pacific Industrial Scale, British Columbia, Canada) to collect the weight shifts on each quadrant (see next section for further details). Following FPSS, TS and 12 QBA attributes were evaluated in the outside testing area, while a single human handler calmly interacted with the calf. The presence of this human handler was intended to facilitate evaluation of the different aspects of these subjective methods. Following Sant’Anna and da Costa (2013), TS is a five point scale, with the neutral value (3) removed, where one indicates calm and five indicates wild or aggressive. The 12 attributes of QBA consist of active, agitated, apathetic, attentive, calm, curious, distressed, fearful, happy, irritated, positively occupied, and relaxed (Sant’Anna and da Costa, 2013), which can be grouped into positive and negative behaviors. Each attribute was scored on a 136 mm line indicating the extent of expression, where the far left and far right refer to no expression and complete expression of the attribute, respectively. Evaluators were given a list of QBA in a specific order, however evaluators scored calves as the attributes were able to be assessed (i.e., based on calf behaviors), therefore a specific order of scoring QBA was never consistent across calves or evaluators. Each calf was measured less than three minutes and then sorted into a holding pen for management purposes.

### FPSS Measurements

The FPSS provides a novel method of quantifying cattle temperament while also weighing the animal (Supplementary Figure S1). The FPSS has scales in each quadrant and connects to a computer controlled by a worker. Prior to the calf entering the scale, the worker first enters the tag number of the calf. Once the calf is on the scale, the worker starts recording weights on the four-platform for at least 45 seconds by starting the recording software. The FPSS computer software is able to record approximately 15 records per second. The worker also keeps a log of any issues encountered with the calf, large movements, and where those issues fall in the records, however these do not influence the selection of records used. Following data collection, FPSS records of each animal were reviewed for quality before used in subsequent analyses. To do this, the ideal start point for a given animal’s scale records (i.e., when the animal is considered as completely standing on the scale) was identified following the protocol in Figure 1. Once the start point was identified, that point and 499 subsequent records were used to calculate the mean and standard deviation of the total weight. The standard deviation of FPSS measurements (SSD) and the coefficient of variation of the SSD (CVSSD = SSD divided by mean) were used as temperament scores for subsequent analyses. The basis of SSD is that animals that are more temperamental will move more often and have larger standard deviations. The CVSSD was calculated as there was concern the actual weight of the animal would bias the SSD as larger animals may naturally have larger standard deviations in records.

**Figure 1:**
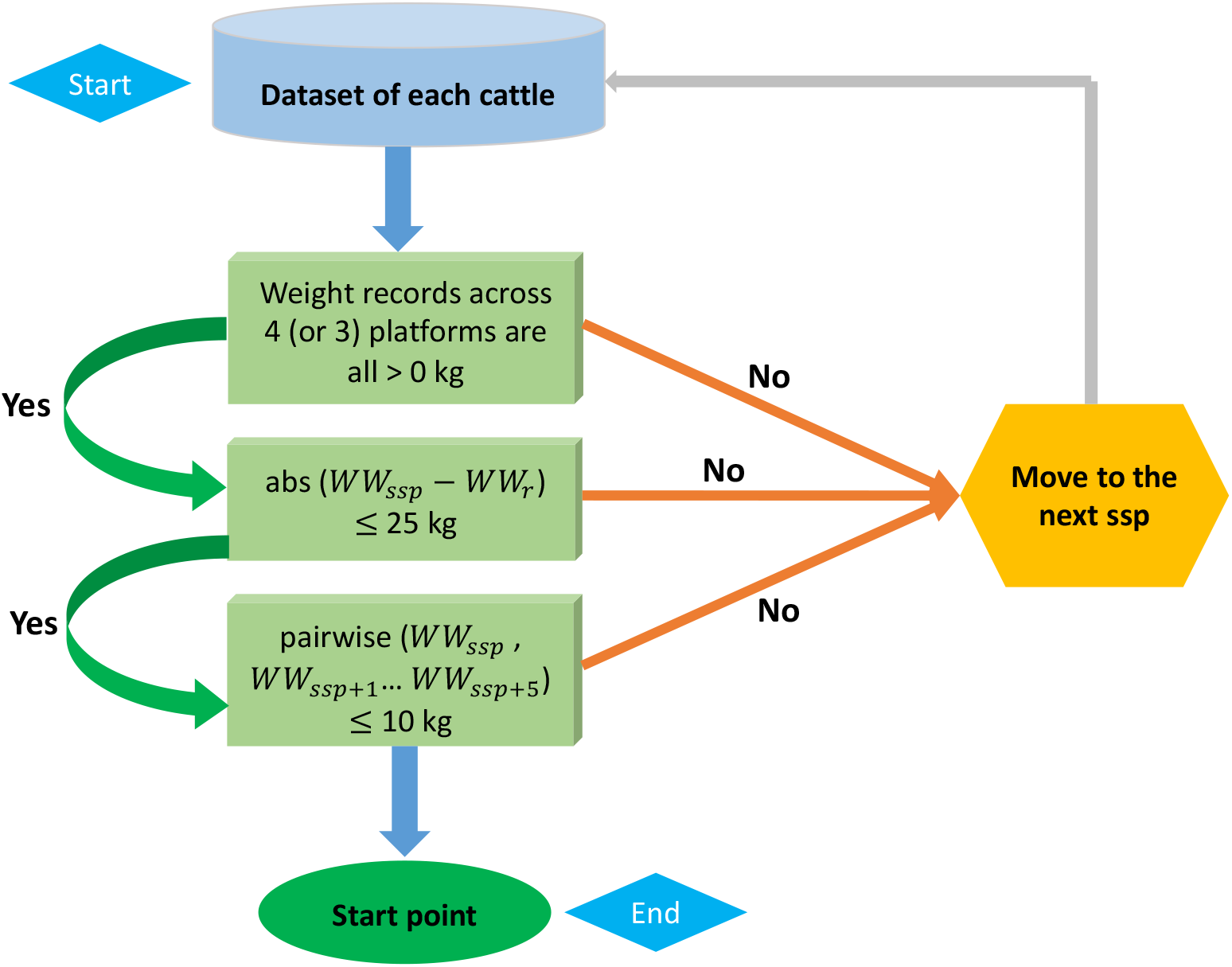
A flow of criteria to identify the start point of four-platform standing scale measurement. The abs and pairwise refer to the absolute difference and pairwise absolute difference, respectively. *WW_ssp_* denotes the weaning weight at current suspected start point (ssp) and *WW*_*ssp*+*i*_ is the weaning weight at the following *i*th point of ssp, where *i* = 1 to 5. *WW_r_* is weaning weight recorded in chute system.

### Exploratory factor analysis

We fitted EFA using subjective measurements including TS and 12 QBA attributes (t = 13). The logic of using EFA is to discover the underlying latent variables or factors (q) to represent observed measurements. Thereby, a network structure between latent variables and phenotypes was first explored and further used for downstream analysis. An EFA model is given as a function of latent factor scores

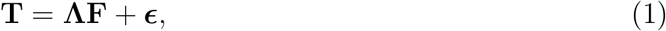

where **T** is a *t* × *n* phenotypic matrix, **Λ** is the *t* × *q* factor loading matrix, **F** is the *q* × *n* latent factor scores, and ***ϵ*** is the *t* × *n* matrix of specific effects. The matrix **Λ** includes the factor loading coefficients, which can be considered as regression coefficients reflecting the relationships between observed phenotypes and underlying latent variables. The variance-covariance structure of **T** is

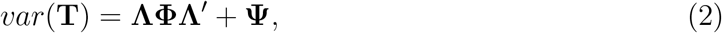

where **Φ** is the variance of factor scores and **Ψ** is the variance of specific effects. With the assumption of 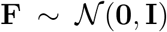, a vector of phenotypes follows a multivariate Gaussian distribution 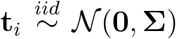 where *i* refers to *i*th individual and **Σ** = **ΛΛ**′ + **Ψ**. The log-likelihood of the factor analysis model is

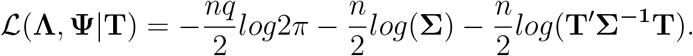

The number of underlying latent variables *q* was determined using a parallel analysis (Horn, 1965). In brief, the eigenvalues of the observed data and simulated data conditioned on the observed data were computed to extract latent variables until the observed data had a smaller eigenvalue than the simulated data. Parameters **Λ** and **Ψ** were estimated by maximizing the log-likelihood of 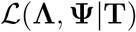 using an iteration method. We used the R package psych (Revelle, 2018) to fit EFA. We posited that DS may not align with other subjective measures since it is collected in a confined setting, whereas other subjective measures used in this study were in a pen with free movement. An additional EFA was fitted including DS to confirm this assumption.

### Bayesian confirmatory factor analysis

Using results from the EFA as a prior, a confirmatory factor analysis (CFA) under the Bayesian framework was fitted following the procedure described in Yu et al. (2019) to obtain factor scores. We assigned the following priors for equations (1) and (2).

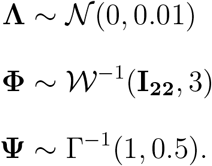

The blavaan R package (Merkle and Rosseel, 2018) coupled with the rstan R package (Carpenter et al., 2017) were applied to solve the Bayesian CFA model. A Markov chain Monte Carlo (MCMC) with 6,000 samples and 3,000 burn-in was adapted to infer the model parameters and in total three MCMC chains were sampled. The model convergence was validated using the combination of trace plots and a potential scale reduction factor (PSRF) less than 1.2 (Brooks and Gelman, 1998). A PSRF compares the estimated variances across chains and within the chain, where a large difference indicates additional Gibbs samplings may be required. This was followed by calculating the factor scores (**F**) of latent variables using the Gibbs samples. When factor scores are considered missing, the **F** is sampled from a conditional distribution of *p*(**F**|***θ***, **T**) (Lee and Song, 2012) using a data augmentation (Tanner and Wong, 1987), where ***θ*** refers to the unknown parameters **Λ**, **Φ**, and **Ψ**. The factor scores of latent variables were summarized from the posterior mean of **F** and considered as new phenotypes in the downstream analysis.

### Bayesian multivariate best linear unbiased prediction

We used a pedigree-based Bayesian multivariate best linear unbiased prediction model to perform genetic analysis of SSD, CVSSD, DS, and latent variables.

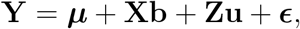

where **Y** is a vector of factor scores with individuals ordered within traits, **X** is the incidence matrix of fixed effects, **Z** is the incidence matrix relating individuals with additive genetic effects, ***μ*** is the vector of intercept, **b** is the vector of fixed effects, **u** is the vector of additive genetic effects, and ***ϵ*** is the vector of residuals. The incidence matrix **X** included birth year by date of collection (*n* = 2 per year), primary breed, and sex following recommendations by (Burrow, 2001; Hulsman Hanna et al., 2019). The joint distribution of **u** and ***ϵ*** follows a multivariate normal

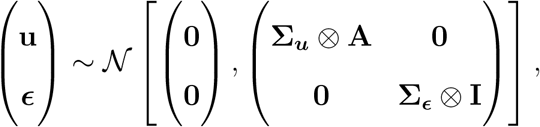

where **A** refers to the numerator relationship matrix, **I** is an identity matrix, **Σ**_***u***_ and **Σ**_***ϵ***_ are genetic and residual variance-covariance matrices, respectively. Flat priors were assigned to ***μ*** and **b**. Inverse Wishart distributions with identity scale matrix and 5 degrees of freedom were assigned for **Σ**_***u***_ and **Σ**_***e***_. The MTM R package (https://github.com/QuantGen/MTM) was employed to infer the parameters in the Bayesian multi-trait linear mixed model and obtain the posterior distribution of these parameters. This was followed by estimating genetic correlations and heritabilities using posterior mean estimates from 10,000 Gibbs samples with 3,000 burn-in and a thinning rate of 5. The model convergence was checked using the trace plots.

## Results

### Phenotypic correlation

The phenotypic correlations between all subjective and objective measurements are displayed in Figure 2. The subjective measurement TS showed a positive correlation with active, fearful, agitated, irritated, and distressed, whereas, a negative correlation with relaxed, calm, and apathetic was found. Among 12 QBA attributes, we observed positive correlations between similar attributes of temperament (e.g., relaxed and calm) and negative correlations for opposite aspects of temperament (e.g., fearful and calm), which supports the use of EFA for this dataset.

**Figure 2:**
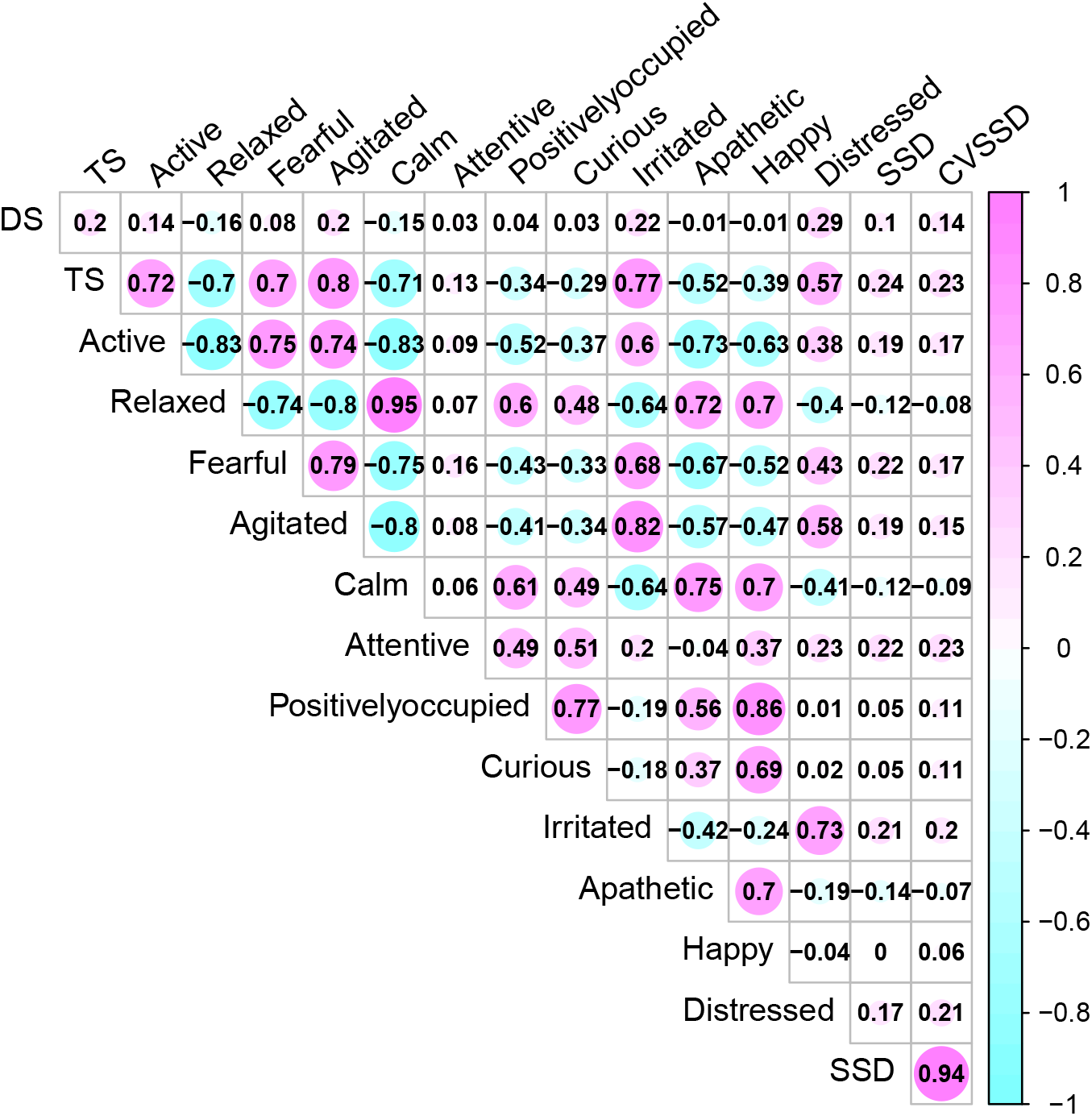
Phenotypic Pearson correlation coefficients between temperament measurements including temperament score (TS), docility score (DS), 12 qualitative behavior assessment attributes, and movement-based scores using four-platform standing scale standard deviation (SSD) and its coefficient of variation (CVSSD).

### Latent structure

The parallel analysis scree plot discovered the first two factors as latent groups (Supplementary Figure S2). The EFA loadings in Figure 3 further identified these two latent groups can be interpreted as *difficult* (factor 1) and *easy* (factor 2) due to loading values. According to Figure 4, the descriptors *difficult* and *easy* were identified because we observed factor 1 has higher loadings for negative temperament attributes (i.e., TS, active, fearful, agitated, irritated, and distressed) and factor 2 has higher loadings for positive temperament attributes (i.e., relaxed, calm, attentive, positively occupied, curious, apathetic, and happy). The EFA factor loadings including DS is displayed in Supplementary Figure S3. As expected, we found that both factors 1 and 2 have low loadings for DS.

**Figure 3:**
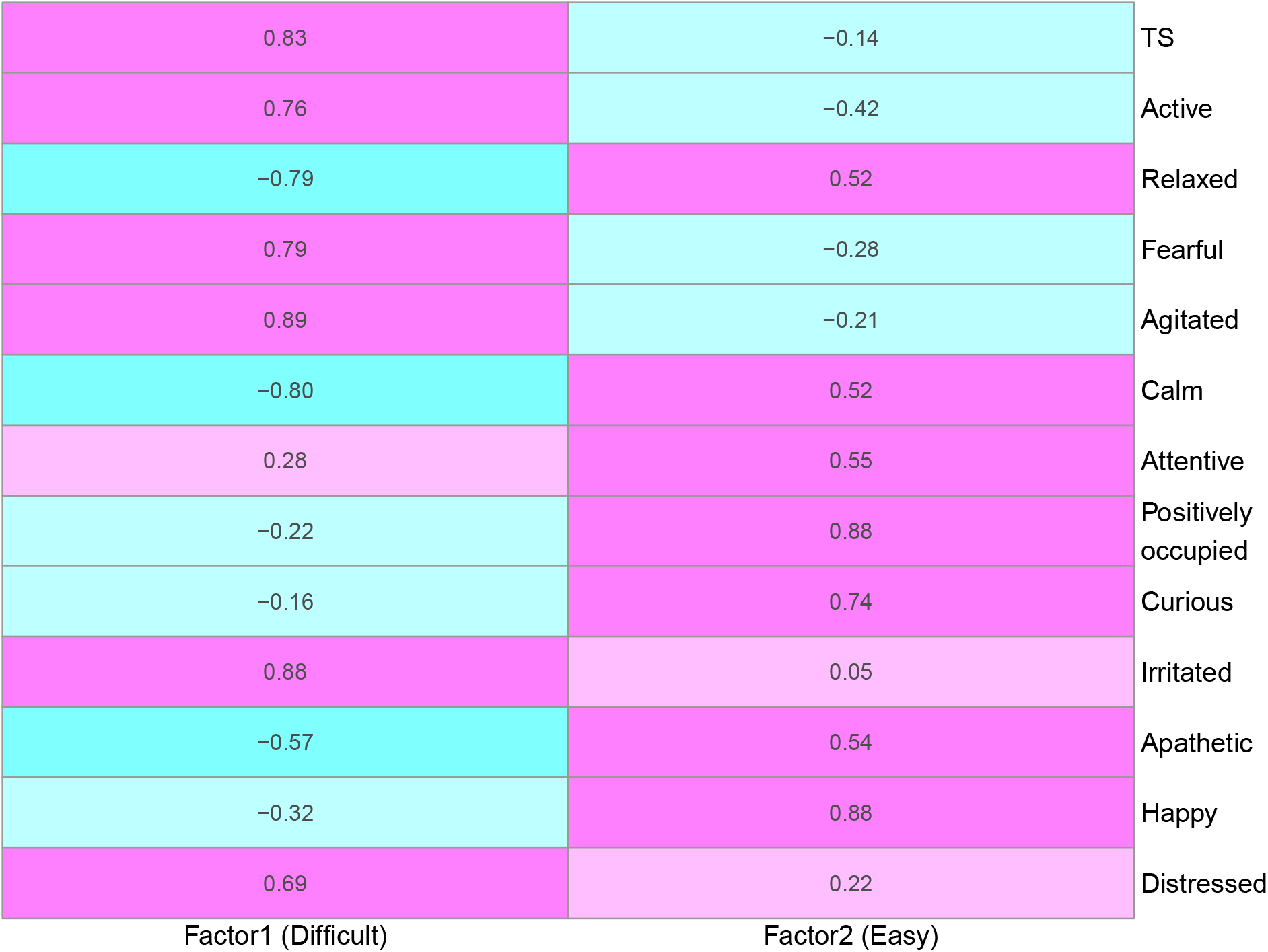
Factor loadings between factors and phenotypes derived from the explanatory factor analysis using temperament score (TS) and 12 qualitative behavior assessment attributes. Positive and negative relationships are denoted as pink and blue, respectively. Factors 1 and 2 are labeled as *Difficult* and *Easy* because of positive loadings on negative and positive temperament attributes, respectively. The degree of shading corresponds to the intensity of the relationships.

**Figure 4:**
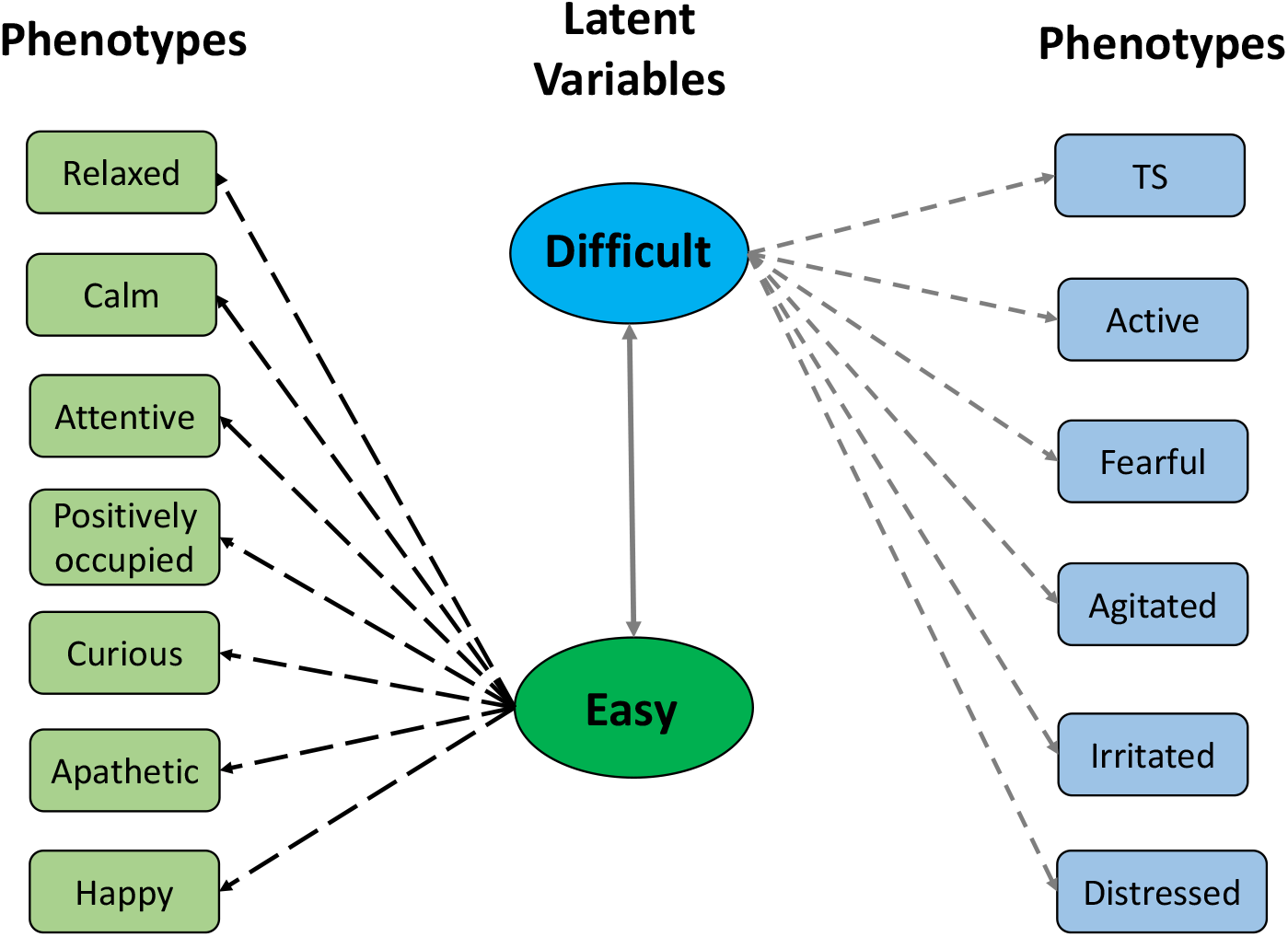
The latent structure between two factors (*Difficult* and *Easy*) and 12 qualitative behavior assessment attributes. The two factors were defined based on their relationships with negative and positive temperament attributes, respectively.

The standardized factor loading coefficients and their posterior standard deviations from CFA assuming latent structure shown in Figure 4 are presented in Table 1. The standardized factor loading coefficients can be interpreted as regression coefficients. Overall, we found two factors have strong contributions to 13 subjective measurements. The factor *difficult* presented a strong positive loading to TS (0.861), active (0.820), fearful (0.840), agitated (0.937), irritated (0.844), and distressed (0.607), suggesting difficult is a comprehensive representation of undesirable aspects of temperament. The factor *easy* showed a positive strong loading to relaxed (0.968), calm (0.982), positively occupied (0.636), curious (0.514), apathetic (0.761), and happy (0.730), indicating an increase of *easy* can result in more desirable temperament.

**Table 1:** Standardized factor loading and corresponding posterior standard deviation from the Bayesian confirmatory factor analysis.

| Latent Variable | Observed Phenotype | Loading | Posterior standard deviation |
| --- | --- | --- | --- |
| Difficult | TS | 0.861 | 0.007 |
| Difficult | Active | 0.820 | 0.010 |
| Difficult | Fearful | 0.840 | 0.008 |
| Difficult | Agitated | 0.937 | 0.004 |
| Difficult | Irritated | 0.844 | 0.009 |
| Difficult | Distressed | 0.607 | 0.016 |
| Easy | Relaxed | 0.968 | 0.002 |
| Easy | Calm | 0.982 | 0.002 |
| Easy | Attentive | 0.079 | 0.025 |
| Easy | Positively occupied | 0.636 | 0.015 |
| Easy | Curious | 0.514 | 0.019 |
| Easy | Apathetic | 0.761 | 0.011 |
| Easy | Happy | 0.730 | 0.012 |

### Genetic relationships among temperament measurements

The inclusion of birth year by date of collection, breed, and sex fixed effects in multivariate analyses of this study follow previous literature. In this study, birth year by date of collection was significant, breed was not significant, and sex was only significant for CVSSD and *easy* based on 95% highest posterior density intervals. The heifers were found to act more temperamental than steers. Effect of sex for these two traits follow trends seen in other literature (Gauly et al., 2001). The estimates of heritability (diagonals) and genetic correlation coefficients (off-diagonals) among SSD, CVSSD, DS, and *easy* and *difficult* factors are shown in Figure 5. The largest negative genetic correlation was observed between *difficult* and *easy* with a posterior standard deviation of 0.02. In reference to DS, SSD, and CVSSD, *difficult* and *easy* had moderate genetic correlations with a posterior standard deviation of 0.16. Intuitively, *difficult* had positive genetic correlations (Figure 5) with DS, SSD, and CVSSD indicating difficulty of handling increased with increasing values of those variables, respectively. Likewise, *easy* had negative genetic correlations with all three variables. The subjective measure of DS showed a positive genetic correlation with objective measures SSD and CVSSD, along with a posterior standard deviation of 0.16 and 0.17, respectively. Thus, a selection on cattle with lower DS measurement may also result in lower SSD and CVSSD measurements. The two objective measurements SSD and CVSSD showed the largest positive genetic correlation with a posterior standard deviation of 0.06. Among all five measurements, we found *difficult* and *easy* showed the largest heritability estimates.

**Figure 5:**
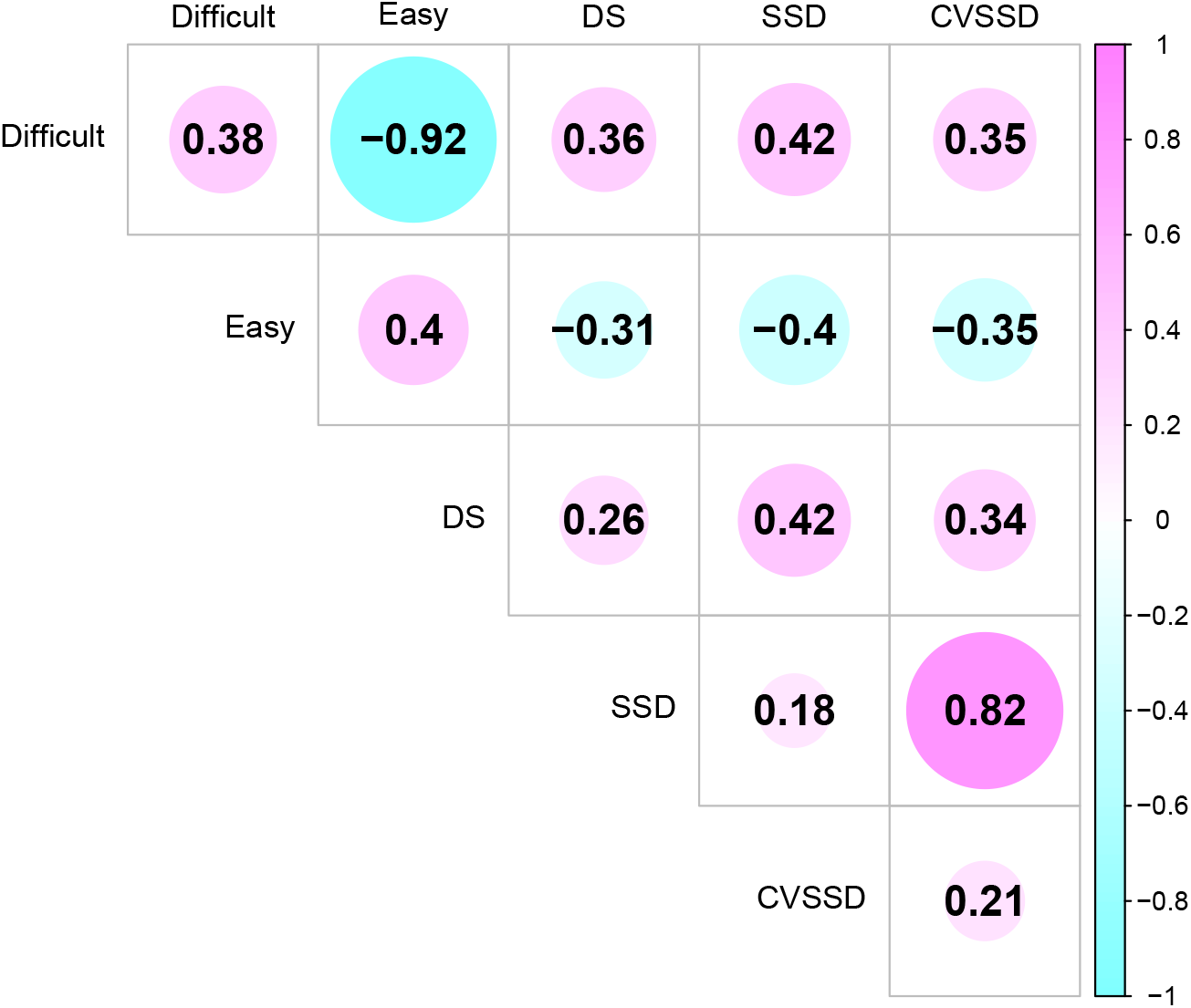
Genetic correlation estimates between latent variables (*Difficult* and *Easy*) and temperament measurements including docility score (DS), standard deviation of total weight on scale over time (SSD), and coefficient of variation of SSD (CVSSD). *Difficult* and *Easy* were defined based on their relationships with negative and positive temperament attributes, respectively. The diagonal elements are the heritability estimates.

The subjective measure DS showed larger heritability estimates than SSD and CVSSD. Heritability estimates of SSD and CVSSD were similar. All heritability estimates had a posterior standard deviation of about 0.06.

## Discussion

### Factor analytic model

Phenotypes are often correlated at the genetic level due to the pleiotropic effect or the linkage disequilibrium among quantitative trait loci. The multivariate modeling has been widely used to model correlated structure by taking the advantage of the genetic or environmental covariance between phenotypes (Henderson and Quaas, 1976; Campbell et al., 2019). The standard multi-trait approach has been proven to be useful for a trait with low heritability or having scarce records (Mrode, 2014). However, it faces a computational challenge when the number of phenotypes included is large. Thus, dimensional reduction methods play an important role in handling high-dimensional phenotypes.

One commonly used approach to study temperament measures is principal component analysis (PCA). This approach calculates principal components (PCs) from a linear combination of observed phenotypes by maximizing the total variance. Napolitano et al. (2012) and Fleming et al. (2013) applied PCA to analyze QBA with the aim of studying dairy buffalo behavior and horse behavior during endurance ride, respectively. Sant’Anna and da Costa (2013) extracted the first principal component from QBA and used it as a new phenotype to study cattle temperament. These studies all suggested traits associated with calm and agitated have a large contribution to the first principle component, which are two extreme characteristics of temperament. The validity of PCs derived from PCA in capturing animal behaviors of both calm and agitated have been supported by the significant correlations with other temperament methods in several studies (Petherick et al., 2002; Sant’Anna and da Costa, 2013). One of the favorable features of QBA is its comprehensive description of temperament by measuring different behaviors. However, the integration of all QBA attributes using PCs with extremely opposite measures (e.g., calm and agitated) may not be desirable because PCA maximizes the total variance, not the variance due to the common signal among measurements. Consequently, selection for temperament based on PCs may be accompanied by substantial risk. Thus, we employed factor analytic modeling for the first time to study temperament measures, which provides a novel approach to investigate multi-phenotypes. The idea behind factor analytic modeling is to represent the observed phenotypes using the unobserved latent variables or factors by maximizing the common variance between correlated phenotypes. When the number of underlying factors are unknown, it is possible to estimate from the data. For instance, de Los Campos and Gianola (2007) performed multi-trait analysis using a factor structure under the Bayesian framework. Alternatively, we can apply a CFA model, when the latent structure is assumed to be known. Peñagaricano et al. (2015) investigated the interrelationships of five latent variables extracted from 19 traits in swine using CFA. Similarly, a Bayesian CFA combined with Bayesian Network was employed to characterize the wide spectrum of 48 rice phenotypes in Yu et al. (2019). These studies determined the latent structure by leveraging the prior biological knowledge between factors and phenotypes. Although the factor analytic model has been applied in animal and plant breeding, there is still paucity of its application to a temperament research.

In this study, we leveraged the combination of EFA and CFA models to identify the mapping between underlying factors and temperament measures, and performed genetic analysis of inferred factors scores. The EFA model aims at estimating the degree of the contributions of factor to phenotypes, while cross-loading (multiple factors contribute to the same phenotype) is allowed. Using the factor loading coefficients, we inferred the latent structure by removing cross-loading. The combination of EFA and CFA modeling relied on a data-driven method to detect the mapping between factors and phenotypes, which is a common case in practice when prior biological knowledge is not available. In this study, we identified two factors *difficult* and *easy* from TS and 12 QBA. This corroborates the findings from previous studies using PCA, where the first principle component has been heavily influenced by both calm and agitated related traits (Napolitano et al., 2012; Fleming et al., 2013; Sant’Anna and da Costa, 2013).

### Temperament measurements

Previous studies reported the estimate of chute score heritability, similar with DS in this study, ranged from 0.11 to 0.34 (Le Neindre et al., 1995; Kadel et al., 2006; Beckman et al., 2007; Hoppe et al., 2010). Burrow and Corbet (2000) suggested objective methods have higher heritabilities than subjective methods of scoring temperament. However, our findings show FPSS measures have slightly lower heritabilities than subjective methods (Figure 5). Haskell et al. (2014) reviewed studies using the objective method of flight speed (exit velocity) and found heritability ranged from 0.05 to 0.70. As suggested by many previous studies, heritability estimates vary in studies based on the population’s phenotypic variation. For temperament studies, this is typically due to differences in experimental design across experiments (e.g., how evaluators interpret and score the animal behavior), different measurement protocols (e.g., the coding of temperament measurements), and breed differences.

Most subjective measures have positive genetic correlations to other subjective measures in previous studies. For example, Grandin (1993) reported a positive correlation between docility test and chute test using Limousin cattle. A positive genetic correlation between race score and crush score (0.530) was detected by Turner et al. (2011) in *Bos taurus* cattle. Sant’Anna and da Costa (2013) discovered a positive genetic correlation between flight speed and temperament index (0.49), which is the first principle component derived from QBA attributes in principle component analysis. An analogous correlation has been detected in this study, where DS showed a positive genetic correlation with *difficult* (0.36), and a negative genetic correlation with *easy* (−0.31) as shown in Figure 5. *Difficult* and *easy* exhibited a large negative genetic correlation (−0.92) which is expected based on the pattern of the temperament measures they loaded to. In this study, subjective methods of DS and *difficult* displayed a moderate positive genetic correlation with objective measurements of SSD (0.42 and 0.42) and CVSSD (0.34 and 0.35). Turner et al. (2011) reported the flight speed has a positive genetic correlation with race score (0.210) and crush score (0.321). Parham et al. (2019) found the exit score and exit velocity capture the same temperament behavior based on a high genetic correlation (0.81). A moderate genetic correlation between flight speed and chute test score has been reported by previous studies (Hoppe et al., 2010; Cafe et al., 2011). These genetic correlations may vary with the differences in breeds, beef production system, evaluator design (e.g., the number of evaluator presented in the study and how evaluator was selected), and/or the number of traits included in a multi-trait analysis. However, genetic correlations found in this study are consistent with previous studies.

DS used in this study, which is also known as chute score (Grandin, 1993), has been widely used in the cattle industry due to its convenience. However, the application of DS is still relying on the human evaluator, which suggests a lack of automation and consistency across populations evaluated (Hieber, 2016; Celestino et al., 2019). Furthermore, DS is the only measurement of temperament with the animal under a restrained condition in contrast with other measurements, which is supported by poor factor loading scores relative to other measures in Supplementary Figure S3. Therefore, we did not combine DS with other subjective methods for EFA. Furthermore, because DS and the two FPSS measures showed similar correlations with *difficult* and *easy* latent variables, the use of FPSS over DS is preferred. This is because the FPSS measures provide automatic, objective, accurate, and consistent measures of temperament rather than relying on evaluator experience and reporting that is needed for DS. It is unlikely, however, that replacement of current scales in cattle production will occur soon due to this. However, the theory of using movement-based scores for temperament has been supported by Sebastian et al. (2011) and Bruno et al. (2018), indicating that replacement of DS with a cost-effective movement-based measure is feasible for genetic selection purposes. Even though DS and FPSS measures identify similar selection on *difficult* and *easy* attributes based on pedigree and correlation coefficients, it is unclear if similar biological pathways or systems are being selected on. Expanding this work to include molecular data to identify genomic relationships among animals and their temperament scores is needed to clarify if selection using movement-based methods can really replace the use of DS in the cattle industry.

## Conclusions

Temperament in cattle is a complex trait that is currently being collected in the cattle industry using the subjective DS method. Our study proves that 1) DS (a restrained score) does not align with temperament attributes that are not restrained and 2) measures captured using the FPSS (objectives measures) have similar genetic correlations to DS. This provides support that movement-measuring devices such as the FPSS can be a feasible replacement for DS and mirror more non-restrained attributes of cattle temperament. Furthermore, our study is the first to implement EFA, Bayesian CFA, and multivariate factor analytic modeling approaches to identify and confirm suspected relationships among multiple traits captured on the same set of animals. We contend that the multivariate factor analytic model applied to the current cattle temperament study provides a new avenue to unravel the complexity of animal behaviors.

## Data Availability Statement

The datasets used in the current study are available from the corresponding author Lauren L. Hulsman Hanna

## Ethics Statement

All procedures were reviewed and approved by the Institutional Animal Care and Use Committee of North Dakota State University (protocols A15015 and A18005).

## Author contributions

This temperament study was conceived and implemented by LLHH, CRD, SAW, and DGR. EFC performed data entry and audit of raw temperament data and pedigree. HY and GM conceptualized applying factor analytic modeling. HY performed the statistical analysis, initial interpretations of results, and drafted the manuscript. HY, GM, EFC, CRD, SAW, DGR, and LLHH revised the manuscript. All authors read and approved the final manuscript.

## Acknowledgements

The authors wish to express their gratitude to the Central Grasslands Research Extension Center director and staff for their assistance with this project, specifically Bryan Neville, Kevin Sedivec, Stephanie Becker, Jessalyn Bachler, Cody Molle, and Rodney Schmidt. Additionally, data collection for this manuscript would not have been possible without the assistance of Kelsey Amborn, Friederike Baumgaertner, Nayan Bhowmik, Dani Black, Justin Crosswhite, Mellissa Crosswhite, Sarah Cushing, Erin Gaugler, Ashley Giedd, Michael Gonda, Lauren Goetze, Jordan Hieber, Alexis Hoelmer, Logan Hulst, Leah Maertens, Kacie McCarthy, Mikayla Miller, Nicolas Negrin, Blaine Novack, William Ogdahl, Nathan Olson, Felipe da Silva, Andrea Strong, Xin Sun, and Sarah Underdahl. This manuscript has been released as a pre-print at bioRxiv (Yu et al., 2020).

## Supplementary Material

