## Supplementary material for "Deciphering cattle temperament measures derived from a four-platform standing scale using genetic factor analytic modeling"

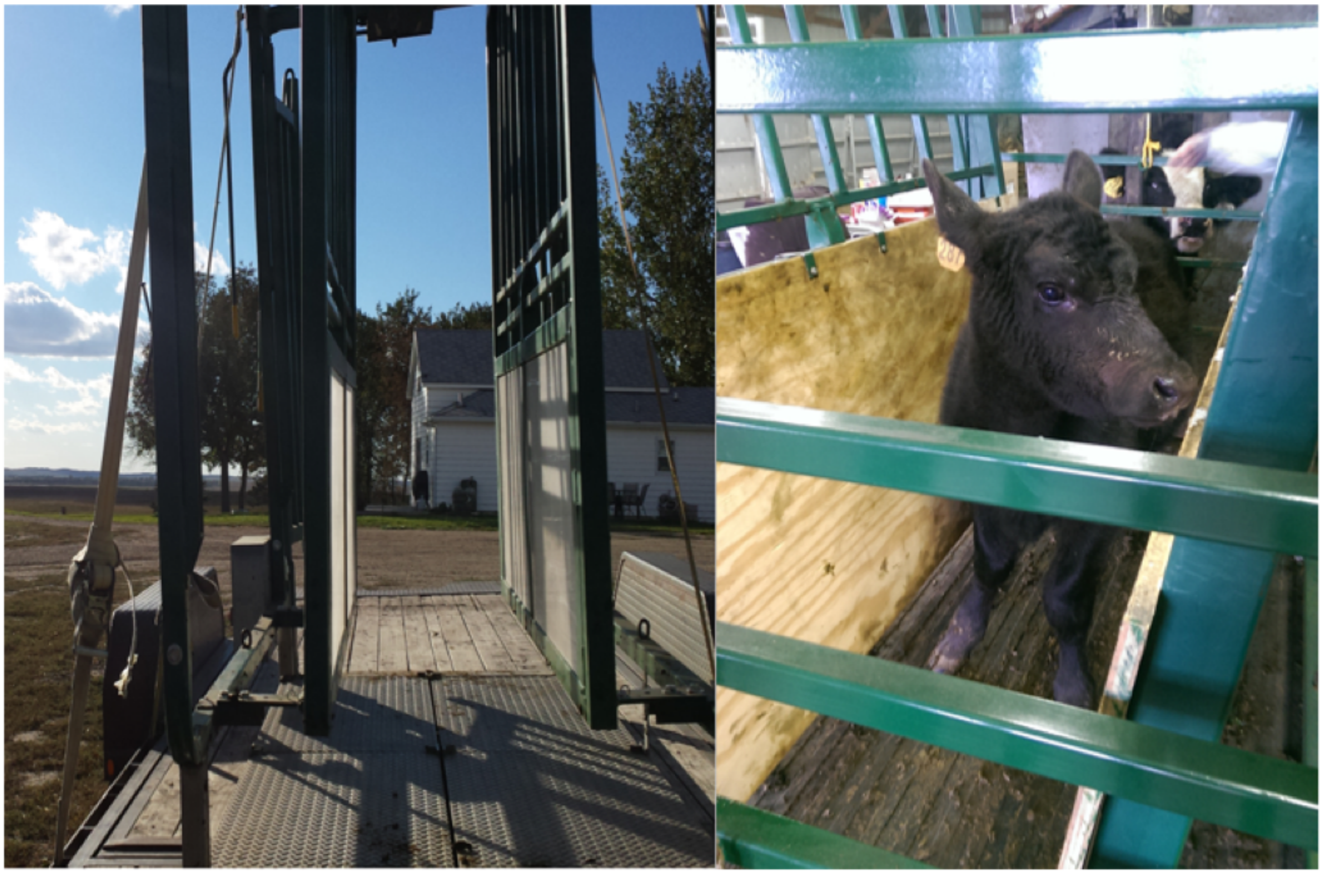

Figure S1: Example of the four-platform standing scale in transport and use during data collection.

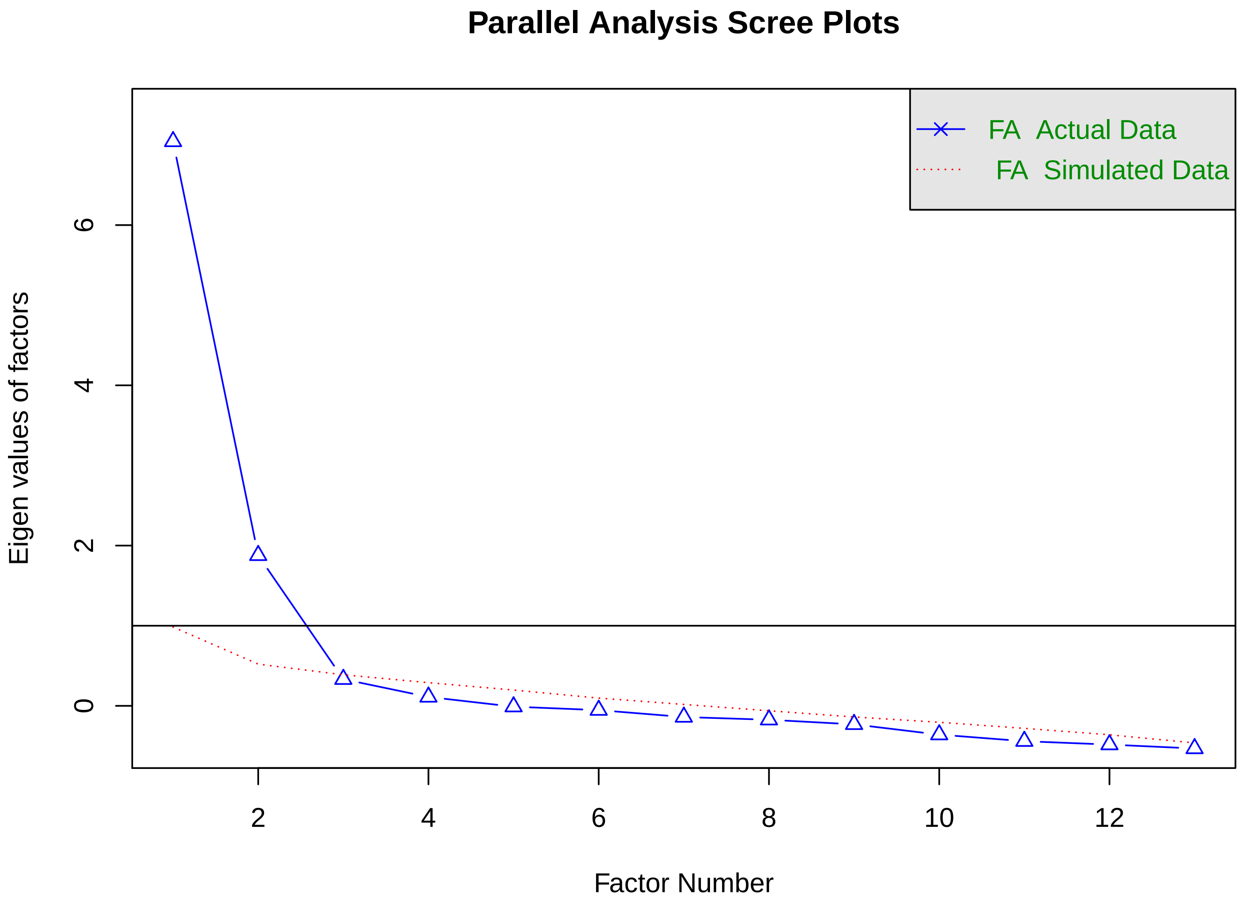


Figure S2: A parallel analysis scree plot to identify the number of factors.

| | Factor1 (Difficult) | Factor2 (Easy) |
|---|---|---|
| DS | 0.26 | 0.06 |
| TS | 0.81 | −0.25 |
| Active | 0.70 | −0.52 |
| Relaxed | −0.72 | 0.62 |
| Fearful | 0.74 | −0.38 |
| Agitated | 0.86 | −0.33 |
| Calm | −0.72 | 0.62 |
| Attentive | 0.35 | 0.51 |
| Positively occupied | −0.10 | 0.90 |
| Curious | −0.06 | 0.75 |
| Irritated | 0.88 | −0.07 |
| Apathetic | −0.49 | 0.61 |
| Happy | −0.20 | 0.91 |
| Distressed | 0.71 | 0.13 |

Figure S3: Factor loadings between factors and phenotypes derived from explanatory factor analysis using temperament score (TS), 12 qualitative behavior assessment attributes, and docility score (DS). Positive and negative relationships are denoted as pink and blue, respectively. Factors 1 and 2 are labeled as *Difficult* and *Easy* because of positive loadings on negative and positive temperament attributes, respectively. The degree of shading corresponds to the intensity of the relationships.

Table S1: Basic means and standard deviations for subjective and objective measures of temperament in weaning age calves group overall, by primary breed, and by sex within primary breed. Primary breed (PB) refers to the calve having 50% or more of Angus (AN) or Hereford (HH) breeding based on known pedigree. Sex is either heifer (H) or steer (S) as bulls were not included in the study. Subjective measures of temperament included docility score (DS; 1 to 6 scale), temperament score (TS, 1 to 5 scale with 3 removed), and 12 qualitative behavior assessment (QBA, 0 to 136 mm line) attributes (grouped and alphabetized by negative and positive characteristics, respectively) of: Active, Agitated, Attentive, Distressed, Fearful, Irritated, Apathetic, Calm, Curious, Happy, Positively occupied, and Relaxed. Objective measures were captured on a four-platform standing scale and included the standard deviation of total weight over a set number of records (SSD) and its coefficient of variation (CVSSD).

| Average | PB | Sex | DS | TS | Active | Agitated | Attentive | Distressed | Fearful | Irritated | Apathetic | Calm | Curious | Happy | Positively occupied | Relaxed | SSD | CVSSD |
|---|---|---|---|---|---|---|---|---|---|---|---|---|---|---|---|---|---|---|
| Overall | | | 1.68± 0.55 | 1.90± 0.78 | 59.20± 26.36 | 28.07± 21.90 | 57.76± 24.33 | 11.89± 11.70 | 23.96± 19.01 | 22.63± 19.47 | 46.36± 28.80 | 73.68± 35.41 | 32.81± 23.77 | 33.91± 28.86 | 34.30± 23.99 | 70.98± 34.05 | 40.23± 23.73 | 0.09± 0.06 |
| | AN | | 1.70± 0.56 | 1.90± 0.77 | 58.70± 26.02 | 27.80± 21.37 | 58.24± 22.98 | 12.30± 11.81 | 23.53± 18.72 | 22.60± 18.93 | 47.45± 29.12 | 74.22± 35.32 | 33.78± 24.08 | 35.30± 29.32 | 35.34± 24.55 | 71.76± 33.87 | 40.25± 23.84 | 0.09± 0.06 |
| | | H | 1.71± 0.58 | 1.92± 0.78 | 60.00± 25.73 | 28.37± 21.46 | 60.23± 23.14 | 13.07± 12.71 | 24.33± 18.47 | 23.35± 19.38 | 46.27± 29.75 | 73.78± 34.33 | 34.3± 23.66 | 35.57± 28.72 | 35.51± 24.54 | 71.01± 33.18 | 39.04± 22.35 | 0.09± 0.06 |
| | | S | 1.70± 0.54 | 1.87± 0.76 | 57.39± 26.31 | 27.3± 21.31 | 56.85± 22.42 | 11.48± 10.66 | 22.86± 19.03 | 21.94± 18.46 | 48.75± 28.61 | 74.91± 36.4 | 33.65± 24.5 | 35.49± 30 | 35.63± 24.59 | 72.71± 34.54 | 41.57± 25.09 | 0.09± 0.06 |
| | HH | | 1.54± 0.45 | 1.90± 0.90 | 62.38± 28.14 | 29.77± 25.12 | 54.17± 32.26 | 8.74± 10.24 | 26.69± 20.32 | 22.62± 22.97 | 38.74± 25 | 70.09± 35.83 | 25.99± 20.12 | 24.08± 23.01 | 27.01± 17.83 | 65.65± 34.91 | 40.01± 23.04 | 0.08± 0.05 |
| | | H | 1.55± 0.43 | 2.00± 0.93 | 65.31± 28.92 | 32.25± 26.82 | 56.45± 40.99 | 10.18± 11.16 | 28.48± 21.54 | 25.83± 24.71 | 35.74± 23.29 | 66.31± 37.16 | 23.34± 18.44 | 23.75± 22.85 | 26.4± 18 | 62.68± 35.96 | 41.27± 24.48 | 0.08± 0.05 |
| | | S | 1.53± 0.46 | 1.83± 0.88 | 60.06± 27.42 | 27.8± 23.63 | 52.37± 23.23 | 7.59± 9.35 | 25.27± 19.28 | 20.09± 21.27 | 41.11± 26.13 | 73.07± 34.63 | 28.09± 21.2 | 24.35± 23.24 | 27.49± 17.76 | 68± 34.05 | 39.02± 21.91 | 0.08± 0.05 |
